# Ventral tegmental area GABA neurons mediate stress-induced anhedonia

**DOI:** 10.1101/2020.05.28.121905

**Authors:** Daniel C. Lowes, Linda A. Chamberlin, Lisa N. Kretsge, Emma S. Holt, Atheir I. Abbas, Alan J. Park, Lyubov Yusufova, Zachary H. Bretton, Ayesha Firdous, Joshua A. Gordon, Alexander Z. Harris

## Abstract

Stressful experiences frequently precede depressive episodes^1^. Depression results in anhedonia, or disrupted reward-seeking, in most patients^2^. In humans^3,4^ and rodents^5,6^, stress can disrupt reward-seeking, providing a potential mechanism by which stress can precipitate depression^7-9^. Yet despite decades investigating how stress modulates dopamine neuron transmission between the ventral tegmental area (VTA) and nucleus accumbens (NAc), the underpinnings of the stress-anhedonia transition remain elusive^10-13^. Here we show that during restraint stress, VTA GABA neurons drive low frequency NAc LFP oscillations, rhythmically modulating NAc firing rates. The strength of these stress-induced NAc oscillations predict the degree of impaired reward-seeking upon release from restraint. Inhibiting VTA GABA neurons disrupts stress-induced NAc oscillations and reverses the effect of stress on reward-seeking. By contrast, mimicking these oscillations with rhythmic VTA GABA stimulation in the absence of stress blunts subsequent reward-seeking. These experiments demonstrate that VTA GABA inputs to the NAc are both necessary and sufficient for stress-induced decreases in reward seeking behavior, elucidating a key circuit-level mechanism underlying stress-induced anhedonia.

## Main

Most patients with major depressive disorder have substantial reward processing impairments^14-16^. Existing therapies incompletely treat anhedonia^17-23^ and its presence strongly predicts clinical outcome^24,25^. Neural activity in the ventral tegmental area (VTA) and nucleus accumbens (NAc) are key to reward processing^26-29^. Yet despite 40 years of research into the dopamine (DA) anhedonia hypothesis^10^, we do not understand how stress disrupts the different aspects of reward processing and its underlying neural circuitry. By contrast, VTA GABA neurons are well situated to mediate the impact of stress on reward-seeking^7^. These neurons fire in response to aversive stimuli^26^ and play a crucial role in calculating reward prediction^27^. They are increasingly recognized to play important roles in stress-mediated addiction behavior^30^, yet previous studies investigating the role of VTA-NAc on stress-induced reward-seeking deficits focused on dopaminergic signaling^12,13,31^. Here, we tested whether VTA GABA projections to the NAc mediate stress-induced anhedonia.

To determine how stress impacts reward processing, we trained mice to collect rewards after hearing a reward-predicting cue (CS+). After a 1.5 second delay from CS+ onset, reward was available for 5 seconds. After reaching stable performance, we recorded VTA-NAc reward circuit activity in mice over two days as they either explored a familiar environment or underwent acute restraint stress in a counterbalanced order (**Fig. 1a**). A prominent low frequency (2-7 Hz) oscillation of local field potential (LFP) activity emerged in the NAc (**Fig. 1b,c; Extended Fig. 1a**) during the restraint stress. A restraint-induced oscillation also appeared in other brain regions, including the prefrontal cortex, where stress has previously been reported to induce low-frequency oscillations^32,33^. However, simultaneous recordings revealed that the restraint-induced oscillation was largest in the NAc (**Extended Fig. 1b,c**). The magnitude of the restraint-induced oscillation did not differ between the core and shell of the NAc (**Extended Fig. 1d**). This restraint-induced oscillation straddles the hippocampal theta frequency range (4-12 Hz), yet simultaneous recordings in the hippocampus and NAc revealed that the restraint-induced oscillation was distinct from hippocampal theta (**Extended Fig. 1e**). We did not observe similar oscillations during periods of voluntary immobility in the familiar environment, suggesting that this neural activity reflects restraint, rather than decreased movement (**Extended Fig. 1f**). As further evidence that stress induces this oscillation, it persisted at the same magnitude throughout the restraint session and only abated when the mice were released from restraint (**Extended Fig. 1g**). Moreover, exposure to another stressor (a mouse aggressor in socially defeated mice) induced a similar low-frequency oscillation in the NAc (**Extended Fig. 1h**). These data support rhythmic low-frequency LFP activity in the NAc as a novel electrophysiological biomarker of acute stress.

**Figure 1.**
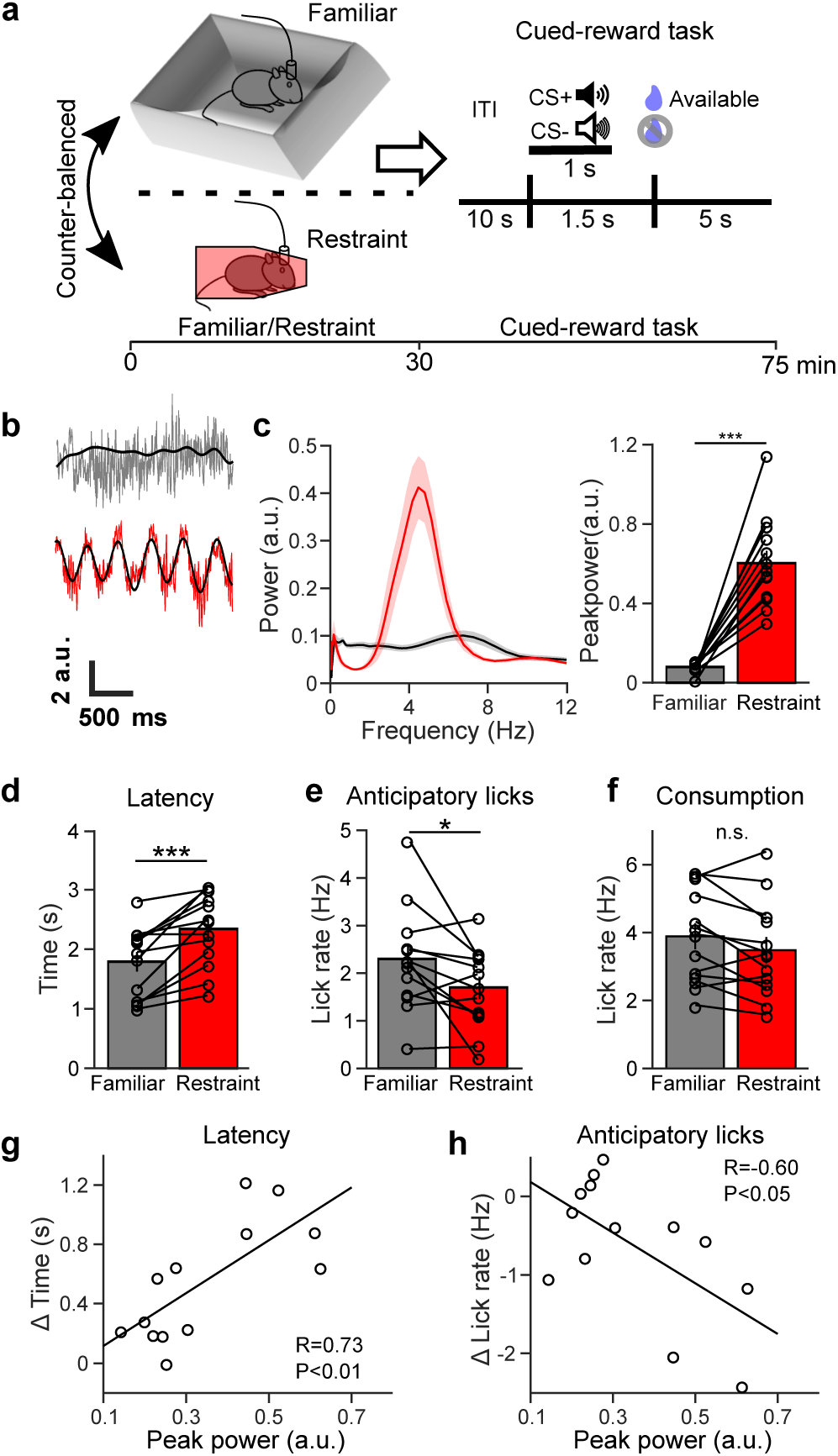
Restraint stress re-organizes the NAc LFP and impairs subsequent reward-seeking. **a.** Experimental design. **b.** Representative traces of the NAc LFP of a mouse freely exploring a familiar environment (grey) or during restraint (red). Black traces are the 2-7 Hz filtered signal. **c.** (Left) Average NAc LFP power spectra of mice exploring a familiar environment (black) or during restraint stress (red). (Right) Peak power in the 2-7 Hz range was higher in restrained mice compared to mice freely exploring a familiar environment (***P<0.001 Wilcoxon signed-rank test W=91, n=13 mice). Data are plotted as mean ± s.e.m. **d.** Latency to reward retrieval in the cued-reward task for mice exploring a familiar environment (black) or experiencing restraint stress (red). Restraint stress increases the time to first lick during the reward availability period (***P<0.001 Wilcoxon signed-rank test, W=89, n=13 mice). Data are plotted as mean ± s.e.m. **e.** Average anticipatory lick rate for mice exploring a familiar environment (black) or experiencing restraint stress (red). Restraint stress decreases average anticipatory lick rate between CS+ onset and reward availability (*P<0.05 paired *t*-test, *t*(12)=3.01, n=13 mice). Data are plotted as mean ± s.e.m. **f.** Average consummatory lick rate for mice exploring a familiar environment (black) or experiencing restraint stress (red). Restraint stress does not affect average lick rate during reward consumption (P>0.05 paired *t*-test *t*(12)=0.935, n=13 mice). Data are plotted as mean ± s.e.m. **g.** Relationship between peak 2-7 Hz NAc power during restraint and the change in latency to reward retrieval from after familiar environment exploration to after restraint (*P<0.01, *r*(11)=0.73, n=13 mice) **h.** Relationship between peak 2-7 Hz NAc power during restraint and the change in anticipatory lick rate from after familiar environment exploration to after restraint (*P<0.05, *r*(11)=-0.60, n=13 mice).

Immediately after undergoing 30 minutes of restraint stress, freely moving mice showed increased latency to retrieve rewards and decreased anticipatory (delay period) licking relative to mice exposed to a neutral environment for an equivalent period (**Fig. 1d,e**). This decreased reward anticipation returned to control levels when measured the following day (**Extended Fig. 1i**). Stress did not significantly alter lick rates during reward consumption (**Fig. 1f**), nor did it decrease general mouse movement (**Extended Fig. 1j**) or non-reward predicting cue (CS-) associated behavior (**Extended Fig. 1k**), suggesting a selective deficit in reward anticipation. Notably, the magnitude of stress-induced oscillations correlated with the degree of reward-seeking impairment (**Fig. 1g,h**), suggesting that this biomarker reflects the circuit activity by which stress disrupts reward-seeking.

Low frequency oscillations typically represent the collective synchronous activity of subthreshold synaptic inputs to a structure^34^. Such input would be expected to modulate NAc neural firing rates. Indeed, restraint stress also robustly affected NAc single unit activity. The firing rates of approximately 75% of NAc single units were significantly modulated by restraint. Most significantly modulated neurons had reduced activity, while a minority had increased firing rates (**Fig. 2a,b**). On average, single unit firing rates became rhythmically entrained to the restraint-induced oscillation (**Extended Fig. 2a,b**). By contrast, prefrontal cortex single units did not phase-lock to the oscillation recorded in the prefrontal cortex (**Extended Fig. 2c**), suggesting that the oscillation originates in the NAc and may be volume conducted to other brain regions. Interestingly, while NAc single units with significantly decreased firing rates showed an increase in phase locking, those with significantly increased firing rates did not, suggesting that the stress-induced oscillation reflects a net inhibitory input to the NAc (**Fig. 2c,d**). Since the magnitude of the oscillation correlates with the degree of impaired reward seeking (**Fig. 1g,h**), these results raise the possibility that an inhibitory input to the NAc may mediate stress-induced reward impairments.

**Figure 2.**
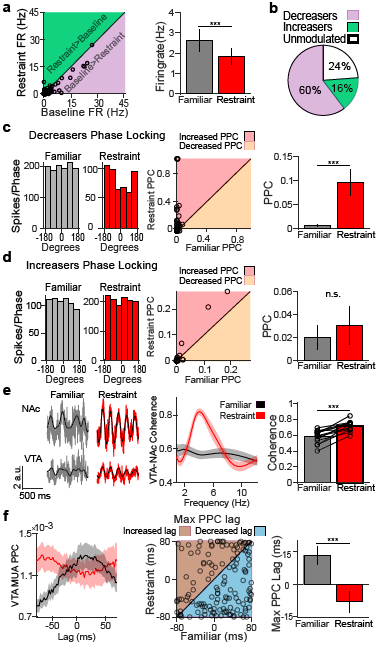
VTA neural activity leads NAc activity during restraint. **a.** (Left) Relationship between the firing rate of NAc single units during familiar environment exploration and restraint stress, with line of equality for reference. Areas above and below the line of equality are shaded green or purple to clarify whether firing rate values increased or decreased (respectively) during restraint. (Right) Restraint reduced the firing rate of NAc single units (**P<0.001 Wilcoxon signed-rank test W=-4537, n=126 neurons). Data are plotted as mean ± s.e.m. **b.** Distribution of NAc neurons excited, inhibited, and unmodulated by restraint (total neurons n=126). **c.** (Left) An example of the spike-phase relationship during familiar environment exploration and restraint for a restraint-inhibited NAc neuron. Activity was uniformly distributed across phase angles in the 2-7 Hz filtered NAc LFP during familiar environment exploration (left), but phase-locked during restraint (right). (Centre) Relationship between the pairwise phase consistency (PPC) of restraint-inhibited NAc neurons during familiar environment exploration and restraint stress, with line of equality for reference. Positive and negative differences are respectively shaded light red and light orange for clarity. (Right) NAc neurons whose firing rates decreased during restraint showed increased phase-locking to the 2-7 Hz filtered NAc LFP during restraint (***P<0.001, Wilcoxon signed-rank test W=2267, n=76 neurons, below left). Data are plotted as mean ± s.e.m. **d.** Same as **c**, but with NAc single units significantly excited by restraint. NAc neurons whose firing rates increased during restraint did not increase phase locking. (P>0.05, Wilcoxon signed-rank test W=26, n=20 neurons). Positive and negative differences are respectively shaded light red and light orange for clarity. Bar graph is plotted as mean ± s.e.m. **e.** (Left) Representative traces of the NAc and VTA LFPs of a mouse freely exploring a familiar environment (grey) or during restraint (red). Black traces are the 2-7 Hz filtered signal. (Centre) Coherence of VTA and NAc LFPs during familiar environment exploration and restraint stress. (Right) Average VTA-NAc coherence in the 2-7 Hz range increases during restraint stress (***P<0.001 paired *t*-test *t*(12)= −6.84, n=13 mice). Line and bar graphs are plotted as mean ± s.e.m. **f.** (Left) Lag analysis of VTA multi-unit activity during familiar environment exploration and restraint stress. Restraint increased synchrony of NAc 2-7 Hz phase with past VTA MUA activity (***P<0.001 Wilcoxon rank-sum test, n_Familiar_=138 multi-units, n_Restraint_=139 multi-units). (Centre) Relationship between lag of max PPC between VTA MUA activity and the 2-7 Hz filtered NAc LFP during familiar environment exploration and restraint, with line of equality for reference. Positive and negative differences are respectively shaded brown and turquoise for clarity. (Right) During restraint VTA MUA activity reorganized from predominantly lagging NAc phase activity to predominantly leading NAc phase activity (***P<0.001 Wilcoxon signed-rank test, W=-3309, n=139 multi-units). Line and bar graphs are plotted as mean ± s.e.m.

We hypothesized that since the VTA projects densely to the NAc and provides GABAergic in addition to dopaminergic input, it could be the source of stress-induced oscillations. Consistent with this hypothesis, coherence between VTA and NAc LFPs increased sharply during restraint stress (**Fig. 2e**) in the same frequency range as the restraint-induced oscillation. This increase in VTA-NAc coherence was significantly greater than any changes in NAc coherence with the prefrontal cortex, amygdala, or hippocampus (**Extended Fig. 2d**). Since the VTA and NAc reciprocally communicate, we inferred directionality by examining the lag at which the phase locking of VTA multiunit activity (MUA) to the NAc restraint-induced oscillation peaks^35,36^. In the familiar environment, VTA MUA activity of the future was best phase-locked to the NAc LFP. By contrast, during restraint stress, VTA MUA activity of the past was best phase-locked to the NAc LFP. These results suggest that during restraint stress the predominant directionality of reward circuit activity is from the VTA to the NAc (**Fig. 2f**).

We next examined the role of VTA inputs in driving the restraint-induced NAc oscillation. We infused the GABA^A^ agonist muscimol (0.5 µg/side) or saline control into the VTA while recording NAc LFP activity in restrained mice. Muscimol treatment significantly reduced the restraint-induced oscillation, suggesting that VTA activity is necessary for NAc restraint-induced activity (**Fig. 3a**). While recent work has proposed that respiration-driven oscillations originating in the olfactory bulbs and piriform cortex represent the true source of NAc oscillations^37,38^, these results and our data showing that the oscillation modulates NAc single unit activity (**Fig. 2c,d**), indicate that restraint-induced oscillations reflect a true NAc oscillation.

**Figure 3.**
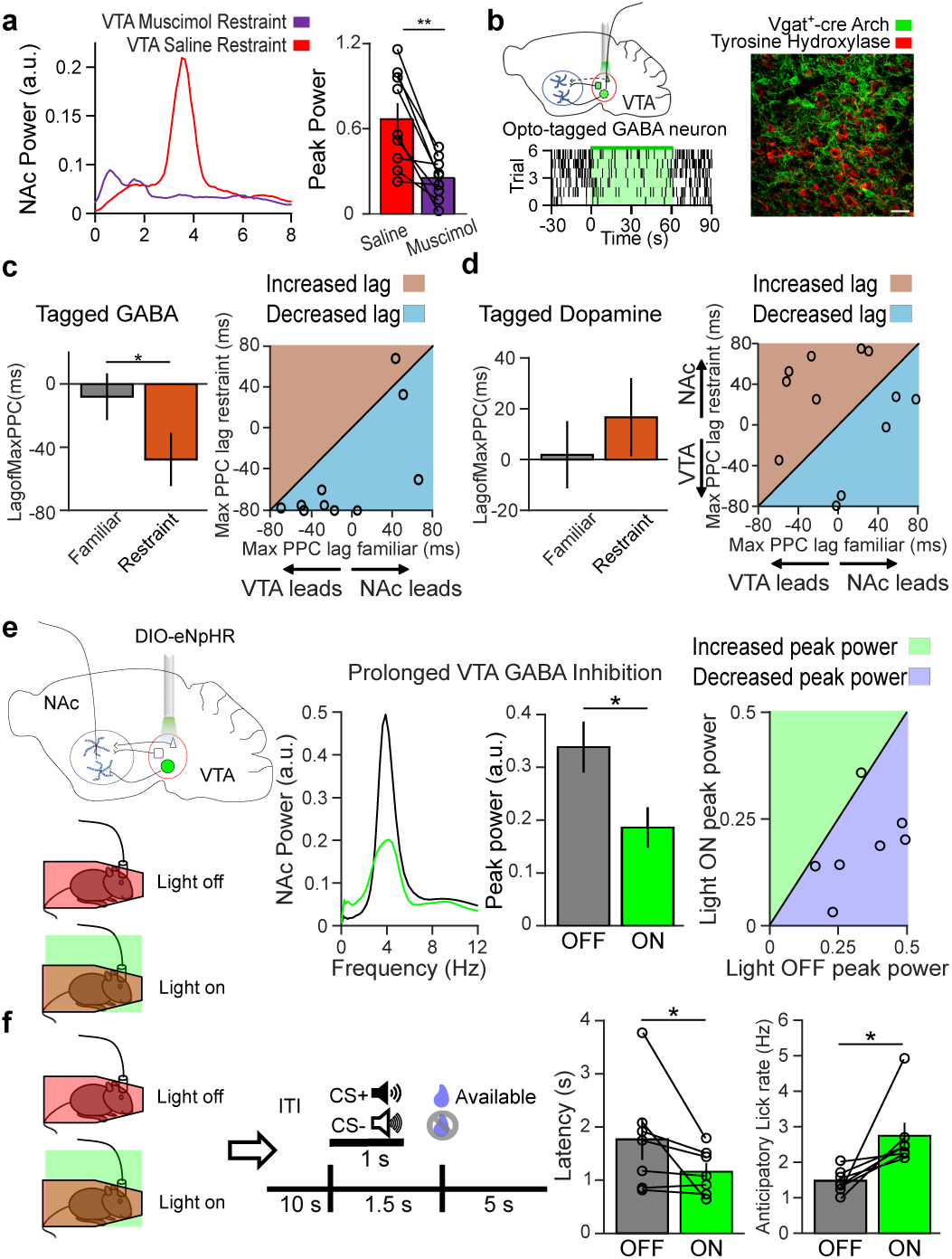
VTA GABA activity contributes to the restraint-induced NAc oscillation and stress-induced reward seeking deficits. **a.** (Left) Example NAc LFP power spectra following infusion of saline (red) or muscimol (violet) into the VTA. (Right) Muscimol infusion reduced the maximum NAc LFP power in the 2-7 Hz range during restraint (**P<0.01 Wilcoxon signed-rank test W=-45, n=9 mice). Bar graph is plotted as mean ± s.e.m. **b.** (Top left) Experimental design. (Bottom left) Example raster plot of opto-tagged GABA neuron. (Right) Example immunofluorescent image of Arch expression (green) in the VTA of a *Vgat*-ires-Cre mouse. Tyrosine hydroxylase counterstain (red) shows expression in non-dopamine neurons. Scale bar is 30 µm. **c.** (Left) Relationship between lag of max PPC between tagged VTA GABA neurons and the 2-7 Hz filtered NAc LFP during familiar environment exploration and restraint, with line of equality for reference. Positive and negative differences are respectively shaded brown and turquoise for clarity. (Right) During restraint the 2-7 Hz filtered NAc LFP became significantly phase-locked with past VTA GABA neuron activity (*P<0.05 Wilcoxon signed-rank test W=-48, n=10 neurons). Data are plotted as mean ± s.e.m. **d.** The same as **c**, but for tagged VTA dopamine neurons. There was no change in the direction of phase-locking between the 2-7 Hz filtered NAc LFP and tagged VTA dopamine neurons (P>0.05, paired t-test *t*(11)=0.754, n=12 neurons). Positive and negative differences were respectively shaded brown and turquoise for clarity. Bar graph is plotted as mean ± s.e.m. **e.** (Left) Experimental design. (Centre) Representative spectrum of NAc LFP during restraint in light off (black) and light on (green) sessions. (Right) Relationship between 2-7 Hz NAc peak power during light off and light on sessions, with line of equality for reference. Positive and negative differences are respectively shaded light green and light blue for clarity. (Far right) Prolonged eNpHR inhibition of VTA GABA neurons reduced the power of the NAc oscillation (*P<0.05 paired *t*-test *t*(6)=3.44, n=7 mice). Data are plotted as mean ± s.e.m. **f.** (Left) Experimental design. (Right) Prolonged eNpHR inhibition of VTA GABA neurons during restraint improved latency to retrieve rewards during the reward availability period (*P<0.05 Wilcoxon signed-rank test W=-26, n=7 mice), as well as anticipatory lick rate between CS+ onset and reward availability (*P<0.05 Wilcoxon signed-rank test W=28, n=7 mice). Data are plotted as mean ± s.e.m.

To assess which VTA cell types provide input to the NAc during restraint stress, we injected Cre-dependent adeno-associated virus (AAV) carrying archaerhodopsin (Arch) into the VTA of *Dat*-ires-Cre and *Vgat*-ires-Cre mice to respectively tag DA and GABA neurons (**Fig. 3b**)^39^. Restraint stress reduced the firing rates of both VTA DA and GABA neurons (**Extended Fig. 3a-c**). However, the lag of peak phase-locking of VTA GABA neural firing was shifted to the past during restraint, indicating that VTA GABA neuron activity precedes the NAc restraint-induced oscillation (**Fig. 3c**). In contrast, the lag of peak phase-locking of VTA DA neural firing did not suggest a particular directionality and did not change with restraint (**Fig. 3d**). Based on these findings, we hypothesized that VTA GABA, but not DA, neurons induce rhythmic NAc activity during restraint stress. To test this hypothesis, we inhibited VTA DA or GABA neurons with archaerhodopsin and compared the power of NAc restraint-induced oscillations. Only inhibiting VTA GABA neurons reduced NAc restraint-induced oscillations (**Extended Fig. 3d-f**). We observed that Arch inhibition of VTA GABA neurons waned over the course of 1 minute, while halorhodopsin (eNpHR) robustly inhibited VTA GABA neurons for 30 minutes (**Extended Fig. 3g,h**). Thus, to determine the role VTA GABA mediated NAc oscillations play in stress-induced anhedonia, we inhibited VTA GABA neurons throughout restraint stress using eNpHR. As expected, prolonged inhibition of VTA GABA neurons during restraint reduced low frequency NAc oscillations (**Fig. 3e)**, but inhibiting VTA GABA neurons in the familiar environment or with control virus had no impact on NAc oscillations (**Extended Fig. 3f,4a**). Critically, inhibiting VTA GABA neurons during restraint reversed subsequent reward-seeking deficits, decreasing latency to lick for reward and increasing anticipatory licking (**Fig. 3f; Extended Fig. 4b**), suggesting that the activity of VTA GABA neurons during stress is necessary for subsequent disrupted reward-seeking.

To assess if VTA GABA neural activity is sufficient to induce a low frequency NAc oscillation, we injected Cre-dependent AAV carrying channelrhodopsin-2 (ChR2) into the VTA of *Vgat*-ires-Cre mice. Stimulating these VTA neurons at a low frequency (4 Hz) drove the emergence of a low frequency oscillation in the NAc (**Fig. 4a, Extended Fig. 5a**). Next, we tested the impact of stimulating VTA GABA neurons on reward anticipation. Thirty minutes of low frequency VTA GABA neuron stimulation in a familiar environment increased reward latency and decreased anticipatory licking during the subsequent reward seeking session (**Fig. 4b, Extended Fig. 5b**). By contrast, stimulating VTA GABA neurons at 20 Hz did not produce an oscillation in the NAc, nor did it significantly blunt reward-seeking (**Extended Fig. 5c,d**). These results indicate that VTA GABA neurons suffice to induce low frequency rhythmic NAc activity and to reduce subsequent reward-seeking, mimicking the effect of stress on NAc physiology and reward behavior.

**Figure 4.**
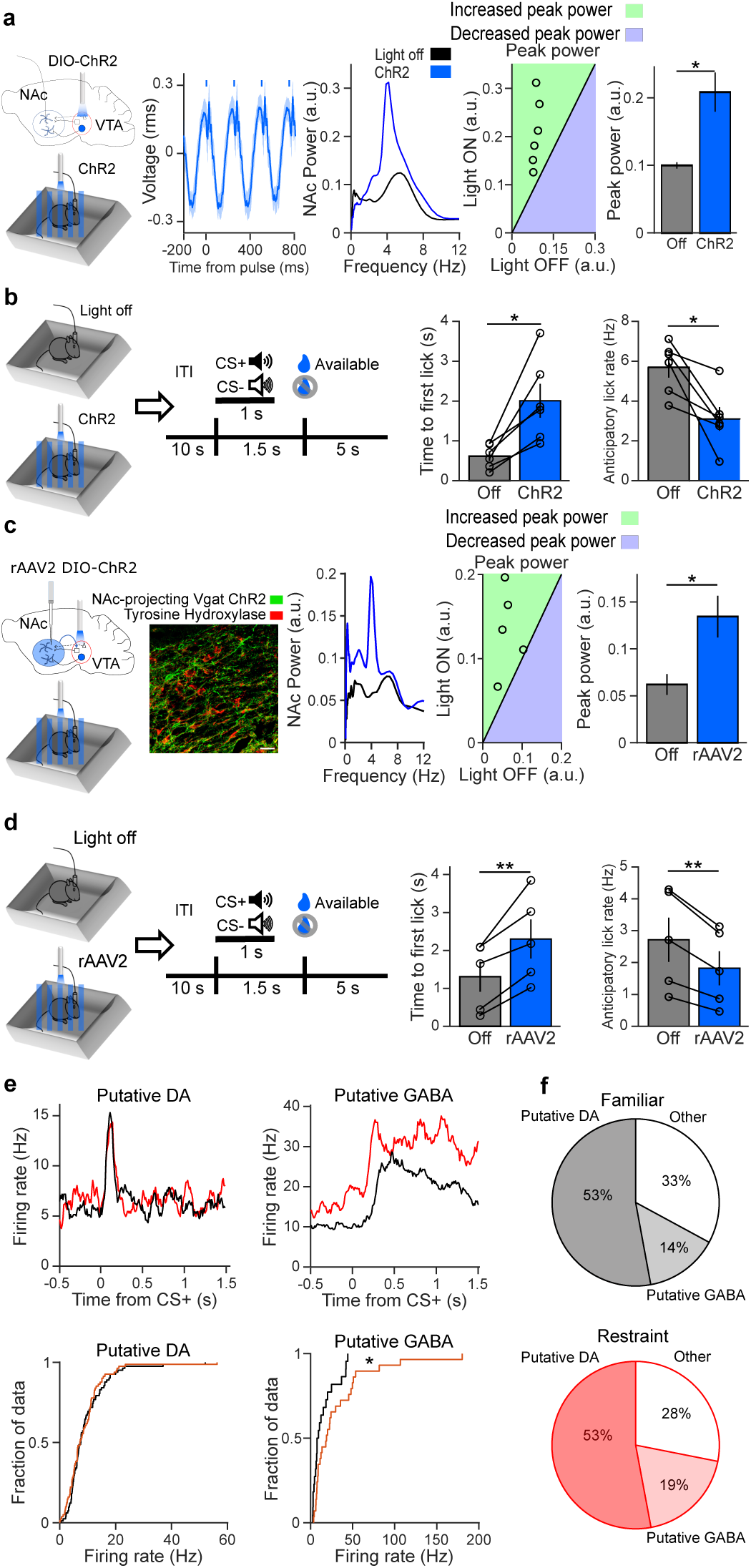
Activation of NAc-projecting VTA GABA neurons produces a low-frequency oscillation and impairs reward-seeking. **a.** (Far left) Experimental design. (Centre left) Event-related potential of the NAc LFP, centered on laser onset (average of n=6 mice, shaded area is s.e.m.). Bars indicate onset and duration of laser pulse. (Centre) Representative NAc spectrum of familiar environment exploration with no stimulation (black, Light off) and with rhythmic low-frequency stimulation (blue, ChR2). (Centre right) Relationship between max 2-7 Hz NAc power during light off and light stimulation, with line of equality for reference. Positive and negative differences are respectively shaded light green and light blue for clarity. (Far right) ChR2 stimulation increased 2-7 Hz NAc peak power (*P<0.05 Wilcoxon signed-rank test W=21, n=6 mice). Data are plotted as mean ± s.e.m. **b.** (Left) Experimental design. (Right) Rhythmic ChR2 stimulation before the cued-reward task impaired latency to reward retrieval (*P<0.05 Wilcoxon signed-rank test W=21, n=6 mice), as well as anticipatory lick rate between CS+ onset and reward availability (*P<0.05 Wilcoxon signed-rank test W=21, n=6 mice). Data are plotted as mean ± s.e.m. **c.** (Far left) Experimental design. (Centre left) Example immunofluorescent image of retrograde ChR2 expression (green) in the VTA of a *Vgat*-ires-Cre mouse. Tyrosine hydroxylase counterstain (red) shows expression in non-dopamine neurons. Scale bar is 30 µm. (Centre) Representative NAc spectrum of familiar environment exploration with no stimulation (black, Light off) and with rhythmic low-frequency stimulation (blue, rAAV2). (Centre right) Relationship between max 2-7 Hz NAc power during light off and light stimulation, with line of equality for reference. Positive and negative differences are respectively shaded light green and light blue for clarity. (Far right) Stimulation of NAc-projecting VTA GABA cell bodies increased 2-7 Hz NAc peak power (*P<0.05 paired *t*-test *t*(4)=-3.00, n=5 mice). Data are plotted as mean ± s.e.m. **d.** (Left) Experimental design. (Right) Rhythmic VTA illumination before the cued-reward task impaired latency to reward retrieval (**P<0.01 paired *t*-test test *t*(4)=-4.73, n=5 mice), as well as anticipatory lick rate between CS+ onset and reward availability (**P<0.01 paired *t*-test *t*(4)=5.40, n=5 mice). Data are plotted as mean ± s.e.m. **e.** (Top) Representative average CS+ evoked firing rates of putative dopamine and GABA neurons in non-stressed (black) and stressed (red) conditions. (Bottom) Distribution of average CS+ evoked firing rates for putative dopamine and putative GABA neurons recorded during familiar environment and restraint stress conditions (total neurons n_Familiar_=155, n_Restraint_=153). Restraint did not affect the CS+ evoked firing rates of putative dopamine neurons (P>0.05, Wilcoxon rank-sum test W=3144, n_Familiar_=82 neurons, n_Restraint_=81 neurons), but enhanced CS+ evoked firing of putative GABA neurons (*P<0.05, Wilcoxon rank-sum test W=203, n_Familiar_=22 neurons, n_Restraint_=29 neurons). **f.** Distribution of putative dopamine and putative GABA neurons recorded during familiar environment and restraint stress conditions (total neurons n_Familiar_=155, n_Restraint_=153).

However, it remained unclear whether direct GABAergic projections from VTA to NAc sufficed to induce low frequency oscillations and blunt reward-seeking. Previous studies reported that stimulating the terminals of NAc-projecting VTA GABA neurons does not impact reward-seeking^40-42^. However, we speculated that robust, rhythmic stimulation mimicking the activity seen during stress might be necessary to reduce reward anticipation. We therefore injected the Cre-dependent retrogradely transported virus rAAV2 carrying channelrhodopsin into the NAc of *Vgat*-ires-Cre mice. Stimulating the VTA cell bodies of NAc-projecting GABA neurons at low frequencies for thirty minutes in a familiar environment induced rhythmic NAc activity and blunted subsequent reward-seeking, with increased reward latency and decreased anticipatory licking (**Fig. 4c,d**), indicating that this small VTA GABAergic sub-population can recapitulate both the physiologic signature of restraint stress and its impact on reward-seeking.

The emergence of VTA GABA neurons as key mediators of stress-induced reward circuit activity suggests a mechanism by which stress disrupts reward-seeking. The orchestrated activity of VTA DA and GABA neurons underlies reward processing. Brief phasic activity of VTA DA neurons in response to reward predicting cues encodes reward anticipation, while prolonged cue evoked firing of VTA GABA neurons sets the level of expectation, resulting in characteristic firing patterns for each cell type^27^ (**Fig. 4e,f**). We reasoned that restraint stress would alter these firing patterns, causing decreased reward anticipation. Indeed, after undergoing restraint mice showed enhanced GABA-like cue-evoked activity, while DA-like cue evoked activity remained intact (**Fig. 4e,f**). These data indicate that stress-induced low-frequency activity of VTA GABA neurons alters their normal function even after stress has abated. Collectively, these data reveal a VTA-NAc circuit mechanism by which stress can cause anhedonia.

## Discussion

Previous efforts to link stress to anhedonia have yielded mixed results^10-13^. Here we show that during acute restraint stress, VTA GABA neurons entrain rhythmic activity in the NAc, producing a newly described 2-7 Hz NAc oscillation. Remarkably, the strength of this oscillation predicts subsequent deficits in reward anticipation. Furthermore, we establish that VTA GABA activity is both necessary and sufficient for stress-induced reward-seeking deficits, as inhibiting VTA GABA neurons during stress reverses the impact on reward-seeking, and mimicking stress-evoked activity in NAc-projecting VTA GABA neurons depresses reward pursuit in unstressed mice.

Our study reveals the cellular mediators of a novel VTA-NAc restraint-induced oscillation. Long-range oscillations form an important motif in neural circuit communication, yet with the exception of theta generation in the medial septo-hippocampal circuit^43^, the cellular mediators of these oscillations remain obscure in most circuits. We show that long-range GABA projections mediate restraint-induced oscillations. NAc-projecting VTA GABA neurons may preferentially inhibit NAc cholinergic interneurons^44^. These cholinergic interneurons represent less than 1% of NAc neurons^45^, yet ramify broadly within the striatum^46^, inhibiting ∼80% of NAc neurons^47^. As a result, VTA GABA neuron rhythmic pacing of NAc cholinergic neurons could underlie the widespread restraint-induced NAc LFP oscillation and single unit firing modulation. Lending further support to this hypothesis, selectively silencing NAc cholinergic interneurons leads to decreased sucrose preference^48^.

While long-range oscillations are typically used to assay the activity of long-range circuits^49^, restraint-induced oscillations seem to reflect the degree to which stress-induced VTA GABA activity induces plastic changes in VTA-NAc circuitry that outlast the stressor and result in impaired reward anticipation. Plasticity affecting VTA GABA neurons has been implicated in the response to cocaine^50^ and stress^7^ and represents an attractive therapeutic target. Our findings thus reveal a VTA GABA-NAc neural circuit which underlies the transition from stress to anhedonia, a critical discovery for advancing depression treatment.

## Figure Legends

<u>Supplemental Figure 1. Characterization of restraint effects on brain-wide LFP organization and reward behavior.</u>

**a.** (Far left) Example NAc spectra from C57BL/6 strain mice. (Centre left) Distribution of frequencies of peak power of restraint-induced oscillation for C57BL/6 mice (n=22 mice). (Centre right) Example NAc spectra from 129/svev strain mice. (Far right) Distribution of frequencies of peak power of restraint-induced oscillation for 129/svev mice (n=19 mice).

**b.** Box-and-whisker plot summarizing the percent change from familiar to restraint in peak 2-7 Hz power of LFP oscillations in the ventral hippocampus (vHPC), basolateral amygdala (BLA), nucleus accumbens (NAc), prefrontal cortex (PFC), ventral tegmental area (VTA) and dorsal hippocampus (dHPC). The percent change of the NAc oscillation was larger than those in the vHPC, BLA, VTA, and dHPC (Kruskal-Wallis Χ^2^=29.1 P<0.0001, df=5; post-hoc Tukey-Kramer comparisons *P<0.05, **P<0.01, ***P<0.001, n_vHPC_=19 mice, n_BLA_=12 mice, n_NAc_=10 mice, n_PFC_=15 mice, n_VTA_=14 mice, n_dHPC_=9 mice). Data are plotted with box-and-whisker plots, which give the median, upper and lower quartiles, and 1.5x interquartile range.

**c.** Simultaneous recordings of the restraint-induced oscillations in the PFC and the NAc revealed that 2-7 Hz peak power was larger in the NAc (*P<0.05, Wilcoxon signed-rank test W=71, n=13 mice). Data are plotted as mean ± s.e.m.

**d.** 2-7 Hz power of the restraint oscillation did not differ between LFP wires targeted to the core and shell of the NAc (P>0.05 Wilcoxon rank-sum test W=58, n_core_=13 mice, n_shell_=10 mice). Data are plotted as mean ± s.e.m.

**e.** Averaged LFP spectra comparing the restraint-induced NAc oscillation to theta oscillations in the dHPC. The restraint-induced NAc oscillation occurs at a lower, distinct frequency band from dHPC theta oscillations (n_NAc_=10 mice, n_dHPC_=9 mice). Data are plotted as mean ± s.e.m.

**f.** (Left) Averaged spectra during periods of immobility during familiar environment exploration (black) and during restraint stress (red). (Right) 2-7 Hz peak NAc power is larger during restraint stress than during immobile periods of familiar environment exploration (***P<0.001 paired *t*-test *t*(21)=10.0, n=22 mice). Data are plotted as mean ± s.e.m.

**g.** Averaged spectrogram of NAc LFP during a pre-restraint baseline, 30 minutes of restraint stress, and immediately after release from restraint (n=11 mice).

**h.** (Left) Representative NAc spectrum of a post-social defeat stress-susceptible mouse during exposure to an empty wire cup (black, Cup) or to a wire cup enclosing a CD1 mouse (red, CD1). (Right) Exposure to a CD1 increased 2-7 Hz NAc peak power in stress-susceptible mice (**P<0.01 Wilcoxon signed-rank test W=45, n=9 mice). Bar graph is plotted as mean ± s.e.m.

**i.** (Left) Latency to reward retrieval for mice exploring a familiar environment on Day 1 (grey) or Day 2 (white) of experiment. Restraint did not have lasting effects on latency to reward retrieval (P>0.05 two-sample *t*-test; n_Day1_=8 mice, n_Day2_=5 mice). (Right) Average anticipatory lick rate for mice exploring a familiar environment on Day 1 (grey) or Day 2 (white) of experiment. Restraint did not have lasting effects on anticipatory lick rate (P>0.05 two-sample *t*-test; n_Day1_=8 mice, n_Day2_=5 mice). Data are plotted as mean ± s.e.m.

**j.** CS+-evoked speed in the cued-reward task. Restraint did not impair mobility (P>0.05 paired t-test *t*(6)=1.53, n=8 mice). Data are plotted as mean ± s.e.m.

**k.** (Left) Average latency to lick during ‘reward’ phase of CS-trials. Restraint stress did not impair lick latency on CS-trials (P>0.05 paired *t*-test *t*(12)=0.813, n=13 mice). (Right) Average anticipatory lick rate during CS-trials. Stress did not affect anticipatory lick rate on CS-trials (P>0.05 Wilcoxon signed-rank test W=-48, n=13 mice). Data are plotted as mean ± s.e.m.

<u>Supplemental Figure 2. Identification of the VTA as possible source of the NAc restraint oscillation.</u>

**a.**An example of an NAc neuron whose activity was uniformly distributed across phase angles in the 2-7 Hz filtered NAc LFP during familiar environment exploration (left), but phase-locked during restraint (right). Positive and negative differences are respectively shaded light red and light orange for clarity.

b. (Left) Relationship between the PPC of all recorded NAc single units during familiar environment exploration and restraint stress, with line of equality for reference. Positive and negative differences are respectively shaded light red and light orange for clarity. (Right) Overall there was an increase in phase-locking between NAc single units and the 2-7 Hz filtered NAc LFP during restraint (***P<0.001, Wilcoxon signed-rank test W=5017, n=126 neurons). Data are plotted with box-and-whisker plots, which give the median, upper and lower quartiles, and 1.5x interquartile range.

**c.** (Left) Relationship between the PPC of PFC single units with the 2-7 Hz filtered NAc LFP during familiar environment exploration and restraint stress, with line of equality for reference. Positive and negative differences are respectively shaded light red and light orange for clarity. (Right) Restraint had no effect on phase-locking strength between PFC single units and the 2-7 Hz NAc LFP (P>0.05 Wilcoxon signed-rank test W=92, n=47 neurons).

**d.** Change in the average 2-7 Hz coherence of the NAc LFP with LFPs located in the vHPC, BLA, PFC, VTA, and dHPC from familiar environment exploration to restraint. The increase in VTA-NAc coherence was larger than the coherence change of all other recorded brain regions (Kruskal-Wallis Χ^2^=32.0 P<0.0001, df=4; post-hoc Tukey-Kramer comparisons ***P<0.001, *P<0.05; n_VHPc_=17 mice, n_BLA_=17 mice, n_PFC_=13 mice, n_VTA_=13 mice, n_dHPC_=7 mice). Data are plotted as mean ± s.e.m.

<u>Supplemental Figure 3. eNpHR is capable of prolonged inhibition of restraint oscillation.</u>

**a.** (Top left) Experimental design. (Top tight) Example immunofluorescent image of Arch expression (green) in the VTA of a *Dat*-ires-Cre mouse. Tyrosine hydroxylase counterstain (red) shows expression in dopamine neurons. (Bottom) Example raster plot of opto-tagged dopamine neuron.

**b.** (Left) Relationship between the firing rate of opto-tagged VTA DA single units during familiar environment exploration and restraint, with line of equality for reference. Positive and negative differences are respectively shaded light green and light purple for clarity. (Right) Restraint inhibited VTA DA firing rate (***P<0.001 Wilcoxon sign-rank test W=-78, n=12 neurons). Data are plotted as mean ± s.e.m.

**c.** (Left) Relationship between the firing rate of opto-tagged VTA GABA single units during familiar environment exploration and restraint, with line of equality for reference. Positive and negative differences are respectively shaded light green and light purple for clarity. (Right) Restraint inhibited VTA GABA firing rate (*P<0.05 Wilcoxon sign-rank test W=-43, n=10 neurons). Data are plotted as mean ± s.e.m.

**d.** (Far Left) Experimental design. (Centre left) Representative spectrum of NAc LFP during restraint in light off (black) and light on (green) epochs. (Centre right) Relationship between 2-7 Hz NAc peak power during light off and light on epochs, with line of equality for reference. Positive and negative differences are respectively shaded light green and light blue for clarity. (Far right) Arch inhibition of VTA dopamine neurons had no effect on the maximum power of the NAc LFP (P>0.05, Wilcoxon signed-rank test W=-7, n=6 mice). Data are plotted as mean ± s.e.m.

**e.** Same as **d.**, but for VTA GABA inhibition. Arch inhibition of VTA GABA neurons reduced the power of the NAc oscillation (*P<0.05 Wilcoxon signed-rank test W=-21, n=6 mice). Positive and negative differences were respectively shaded light green and light blue for clarity. Bar graph is plotted as mean ± s.e.m.

**f.** (Far left) Experimental design. (Centre left) Representative spectrum of NAc LFP during restraint in light off (black) and light on (green) epochs. (Centre right) Relationship between 2-7 Hz NAc peak power during light off and light on epochs, with line of equality for reference. Positive and negative differences are respectively shaded light green and light blue for clarity. (Far right) Illumination of the VTA had no effect on the maximum power of the NAc LFP (P>0.05 paired *t*-test *t*(3)=-2.57, n=4 mice). Bar graph is plotted as mean ± s.e.m.

**g.** (Left) Example traces of efficacy of Arch and eNpHR inhibition of VTA GABA neurons in decreasing the 2-7 Hz NAc oscillation during restraint. (Right) Efficacy of Arch and eNpHR inhibition of VTA GABA neurons in decreasing the 2-7 Hz NAc oscillation during different times in the light on period. Arch inhibition of VTA GABA neurons decreases the 2-7 Hz oscillation during the first 10 s of illumination (*Bonferroni adjusted P<0.05 one-sample *t*-test t(5)=5.21, n=6 mice) but not during the last 10 s of illumination (Bonferroni adjusted P>0.05 one-sample *t*-test t(5)=3.06). By contrast, eNpHR inhibition of VTA GABA neurons successfully decreases the 2-7 Hz restraint oscillation during the first 10 s of illumination (***Bonferroni adjusted P<0.001 one-sample *t*-test *t*(6)=8.72, n=7 mice), 50-60s after illumination onset (*Bonferroni adjusted P<0.05 one-sample *t*-test t(6)=4.18), and 1740-1800 s after illumination onset (**Bonferroni adjusted P<0.01 one-sample *t*-test *t*(6)=5.87). Bar graphs are plotted as mean ± s.e.m.

**h.** (Left) Example traces of the efficacy of Arch and eNpHR inhibition of VTA GABA neurons in decreasing the firing rate of opto-tagged neurons during restraint. (Right) Efficacy of Arch and eNpHR inhibition of VTA GABA neurons in decreasing opto-tagged GABA neuron firing rates during different times in the light on period. Arch significantly reduced the firing rates of tagged GABA neurons 50-60 s after illumination onset (***Bonferroni adjusted P<0.001 one-sample *t*-test t(9)=10.3, n=10 neurons), but not 0-10 s after illumination onset (Bonferroni adjusted P>0.05 one-sample *t*-test t(9)=2.35, n=10 neurons). eNpHR successfully inhibited tagged VTA GABA neurons 0-10 s, 50-60 s, and 1740-1800 s after illumination onset (all ***Bonferroni adjusted P<0.001 one-sample *t*-test; respectively t(7)=15.7, t(7)=13.4, and t(7)=22.0; n=8 neurons). Bar graphs are plotted as mean ± s.e.m.

**i.** (Left) Experimental design. (Centre) Example trace of effect of eNpHR inhibition on 2-7 Hz NAc peak power during familiar environment exploration. (Right) VTA GABA inhibition during familiar environment exploration has no effect on peak 2-7 Hz NAc power (P>0.05 paired t-test t(3)=-0.090, n=4 mice). Bar graphs are plotted as mean ± s.e.m.

<u>Extended Figure 4. eYFP control for prolonged VTA illumination.</u>

**a.** (Far left) Experimental design. (Centre left) Representative spectrum of NAc LFP during restraint in light off (black) and light on (green) sessions. (Centre right) Relationship between 2-7 Hz NAc peak power during light off and light on sessions, with line of equality for reference. Positive and negative differences are respectively shaded light green and light blue for clarity. (Far right) Prolonged VTA illumination during restraint had no effect on peak 2-7 Hz NAc power (P>0.05 paired *t*-test *t*(3)=1.62, n=4 mice). Bar graphs are plotted as mean ± s.e.m.

**b.** (Left) Experimental design. (Right) Prolonged VTA illumination during restraint had no effect on latency to retrieve rewards during the reward availability period (P>0.05 paired t-test t(3)=1.95, n=4 mice), or anticipatory lick rate between CS+ onset and reward availability (P>0.05 paired t-test t(3)=-2.02, n=4 mice). Bar graphs are plotted as mean ± s.e.m.

<u>Extended Figure 5. Rhythmic VTA illumination does not have non-specific effects on Nac physiology or reward-seeking behavior.</u>

**a.** (Far left) Experimental design. (Centre left) Representative NAc spectrum of familiar environment exploration with no stimulation (black, Light off) and with rhythmic low-frequency illumination (blue, eYFP). (Centre right) Relationship between max 2-7 Hz NAc power during light off and rhythmic light illumination, with line of equality for reference. Positive and negative differences are respectively shaded light green and light blue for clarity. (Far right) Rhythmic light illumination had no effect on peak 2-7 Hz NAc LFP power (P>0.05 paired *t*-test *t*(3)= −0.633, n=4 mice). Bar graphs are plotted as mean ± s.e.m.

**b.** (Left) Experimental design. (Right) Rhythmic VTA illumination before the cued-reward task had no effect on latency to retrieve rewards during the reward availability period (P>0.05 paired *t*-test *t*(3)=0.518, n=4 mice), or anticipatory lick rate between CS+ onset and reward availability (P>0.05 paired *t*-test t(3)=-0.982, n=4 mice). Data are plotted as mean ± s.e.m.

**c.** (Far left) Experimental design. (Centre left) Representative NAc spectrum of familiar environment exploration with no stimulation (black, Light off) and with rhythmic high-frequency stimulation (blue, 20 Hz). (Centre right) Relationship between max 2-7 Hz NAc power during light off and rhythmic light illumination, with line of equality for reference. Positive and negative differences are respectively shaded light green and light blue for clarity. (Far right) 20 Hz stimulation did not affect 2-7 Hz NAc peak power (P>0.05 paired *t*-test t(3)= −0.830, n=4 mice). Bar graphs are plotted as mean ± s.e.m.

**d.** (Left) Experimental design. (Right) Rhythmic high-frequency VTA illumination before the cued-reward task had no effect on latency to retrieve rewards during the reward availability period (P>0.05 paired *t*-test t(3)=-2.56, n=4 mice), or anticipatory lick rate between CS+ onset and reward availability (P>0.05 paired *t*-test t(3)=1.51, n=4 mice). Data are plotted as mean ± s.e.m.

## METHODS

### Subjects

4-6-month-old male and female *Vgat*-ires-Cre homozygous mice, *Dat*-ires-Cre heterozygous mice, C57BL/6J mice, and 129/svev mice (The Jackson Laboratory, stock numbers 028862, 006660, 0006664, and 002448 respectively) were used as experimental subjects. Mice were housed with a standard 12 hour light-dark cycle and were given *ad libitum* access to food and water (when not being water restricted for the cued-reward task).

### Surgical Procedures

Mice were anesthetized with 1%–3% vaporized isoflurane in oxygen (1 L/min) and placed in a stereotaxic apparatus. Cre mice were injected with Cre-inducible Archaerhodopsin (AAV5-EF1a-DIO-eArch3.0-EYFP, UNC vector core), Cre-inducible Halorhodopsin (AAV5-Ef1a-DIO eNpHR 3.0-EYFP, Addgene), Cre-inducible Channelrhodopsin-2 (AAV5-Ef1a-DIO hChR2(E123T/T159C)-EYFP, Addgene), or control EYFP virus (AAV5-EF1a-DIO-EYFP, UNC vector core) at two locations within the VTA bilaterally (+/-0.5 ML, −3.4 AP, 4.5 depth below brain surface; 0.5 μL virus per injection. For projection-specific experiments, mice were injected with retrogradely transported Cre-inducible Channelrhodopsin-2 (rAAV2-EF1a-DIO-hChR2(H134R)-EYFP-WPRE-HGHpA, Addgene) at eight locations within the NAc bilaterally (±1.8 mm ML, 0.98 mm AP, −4.12 mm DV; ±0.9 mm ML, 1.5 mm AP, −4.0 mm DV; ±0.6 mm ML, 1.5 mm AP, −3.4 mm DV; ±1.0 mm ML, 1.0 mm AP, −3.4 mm DV; 0.4 μL virus per injection). For microdrive implantation experiments, animals underwent a second surgery to implant the microdrive about 4 weeks after initial viral injection. Animals were again anesthetized and placed in a stereotaxic apparatus. Craniotomies were made to allow for implantation of a bundle of 15 stereotrodes (13 micron tungsten wire, California Fine Wire) and a 200 micron optical fiber in the left VTA (−0.5 ML, −3.4 AP, 4.45 below brain surface); a 200 micron optical fiber in the right VTA; a local field potential (LFP) wire (76 micron tungsten wire, California Fine Wire) in the NAc (1.25 ML, 1.25 AP, 4.0 below brain surface); a ground screw over the cerebellum; and a reference screw over the orbitofrontal cortex. Electrodes were connected to a 32 channel electrode interface board (EIB-36 Narrow, Neuralynx), and the board and wires were fixed to an advanceable custom microdrive. Electrode placements were confirmed by passing current through an electrode at each site (5 mA, 10 s) to generate an electrothermolytic lesion. Mice were anesthetized with ketamine/xylazine prior to generating the lesions and were then perfused. The brains were extracted, cryoprotected, sectioned, mounted, stained, and examined under a microscope to determine lesion placements and characterize eYFP, Arch, eNpHR, and ChR2 expression in VTA.

### Optogenetics

561 nm wavelength light (Opto Engine, LLC), 532 nm wavelength light (OEM laser) or 473 nm wavelength (LaserGlow Technologies) light was delivered at 10 mW via 200 mm diameter, 0.22 NA optical fibers. For Arch experiments, 532 nm light was delivered in 60 second ON/60 second OFF epochs as mice explored a familiar environment and while mice were restrained. For eNpHR experiments, 532 nm or 561 nm light was delivered constantly for 30 minutes while mice were restrained. In addition, 532 nm or 561 nm light was delivered 1 second ON/10 second OFF as mice explored a familiar environment for optogenetic tagging. For ChR2 experiments, mice either received no illumination or 4 Hz stimulation (10 ms pulse width) while exploring a familiar environment. For both stimulation and non-stimulation days, mice received 50 intermittent light pulses at the end of the recording for optogenetic tagging.

### Immunohistochemistry

At the conclusion of experiments (6-8 weeks after viral injection), mice were first anesthetized with a ketamine (100 mg/kg) and xylazine (7 mg/kg) mixture then perfused with 4% paraformaldehyde (Thermo Fisher Scientific), and the brains were extracted and cryoprotected in 30% sucrose in 1X PBS until they sank. 40 µm sections were taken in a cryostat and stored in 1X PBS at 4°C until they were used for immunohistochemistry experiments. The following antibodies and dilutions were used: chicken polyclonal anti-GFP (1:500, Abcam, ab13970), Cy3 donkey anti-sheep (1:200, JacksonImmunoResearch, 713-165-147), sheep polyclonal anti-TH (1:1000, Abcam, ab113), Cy2 donkey anti-chicken (1:200, Jackson ImmunoResearch 703-225-155).

### Cued-reward task

After 5–7 days of post-surgical recovery, the mice were water restricted to 90% of baseline body weight. Prior to experimental recordings, the mice underwent 3 days of habituation to the recording setup: an opto/electrical tether was connected to the head stage preamplifier while the mice explored a small, rectangular, wooden box (24.5 × 34 cm) in 200 lux lighting conditions in 15-20 min daily sessions. Upon reaching their target weight, the mice were trained on a cued-reward task using a custom-built, Arduino-driven rectangular, wooden box in which tones (1 kHz or 5-10 kHz white noise; 1 second duration) produced by a piezo buzzer (Digi-Key) predicted water rewards (15 µL) delivered with a solenoid valve (The Lee Company). The mice were presented with 150 trials of pseudo-randomly interleaved CS+ (predicting reward) or CS-(non-reward predicting). CS+/CS-identity was counterbalanced across mice. After the CS+ ended there was 0.5 second delay, followed by a 5 second period during which reward would be delivered upon detection of a lick at the reward port by an infrared photointerruptor (SparkFun). This reward retrieval period was followed by a 10 second intertrial interval (ITI). Mice were trained until they retrieved rewards on at least 70% of CS+ trials for two consecutive days. Lick detection at the reward port was used to measure the first lick during the reward availability period and the rate at which mice would lick during reward anticipation and reward delivery.

### Restraint

Mice were placed in tapered plastic film restraint bags (Decapicone) punctured with breathing holes before entering 50 mL plastic conical tubes, modified with a slit to allow the headstage and recording tether to pass. The mice used to characterize restraint-induced physiology in Figure 1 and Extended Figure 1 had previously undergone a 5-minute social interaction test not analyzed in this study.

### Social Defeat

CD1 retired breeder males (Charles River laboratories) were used as aggressors. We prescreened CD1 males to ensure that they attacked male mice in under 1 min. For 10 days, we placed the experimental mouse in a different aggressive mouse’s homecage for 10 minutes. The mice then spent the remaining 24 hours across a perforated, transparent divider from the aggressor.

### Social Interaction

Mice explored a chamber (40 cm x 40.5 cm) containing an empty enclosure (10 cm diameter wire cup) for 2.5 min, then explored the same chamber for an additional 2.5 min after we introduced a novel mouse (CD1 male) into the enclosure. We defined the interaction zone using a 7 cm radius around the enclosure. Susceptible mice were defined as those that spent less time interacting with the CD1 mouse than they did with the empty enclosure.

### Pharmacological inactivation

Guide cannulas (26 gauge; Plastics One) were bilaterally implanted into the VTA as described above. An LFP wire was glued to the side of the cannula so that it extended 1.5 mm past the cannula tip. LFP wires were also implanted in the NAc. After recovery from surgery, the mice were habituated to the familiar environment for 3 days. Saline or muscimol (Tocris Bioscience) dissolved in saline (8.8 mM concentration) was microinfused into the VTA while the mice were in the home cage by back loading into a 33-gauge infusion cannula and into polyethylene (PE 20) tubing connected to 1.0 µL Hamilton microsyringe. The infusion cannula protruded 1 mm beyond the guide cannula tip. An infusion volume of 0.13 µL was delivered using a Harvard 11 Plus syringe pump (Harvard Apparatus) at a rate of 0.13 μL/min and the infusion cannula was left in place for 5 minutes post-infusion. After a 20–30 min post-infusion interval, neural activity was recorded as the mice were placed in the familiar environment and then the restraint tube.

### Neural Recordings

A Digital Lynx system (Neuralynx, Bozeman, MT) was used to amplify, band-pass filter (1-1000 Hz for LFPs and 600-6000 Hz for spikes), and digitize the electrode recordings at sampling rates of 2 and 32 kHz for LFPs and spikes, respectively. Single units were clustered based on the first two principal components (peak and energy) from each channel using Klustakwik (Ken Harris) and visualized in Spike Sort 3D (Neuralynx). Clusters were then visually inspected and included or eliminated based on waveform appearance, inter-spike interval distribution, isolation distance, and L-ratio.

### Power, Coherence, Phase-Locking Analysis, and Directionality Analysis

To calculate power, after normalizing the raw LFP data to the root mean square of the voltage signal while the mice were in the familiar environment, the power spectra and coherence of the LFPs were calculated with MATLAB wavelet functions (*cwt* and *wcoherence*). Power and coherence were calculated on 1-minute segments of data and averaged across the entire experimental session. To remove mechanical artifacts and EMG noise, indices with voltage values 3 standard deviations from the mean were omitted when averaging power. To speed-filter data, indices corresponding to when the animal was moving 0-5 cm were included when averaging power. To capture the animal-specific frequency of the restraint-induced oscillation, we measured the peak restraint-induced signal within a 2-7 Hz range and compared it with the corresponding signal in the familiar environment.

For phase locking analysis, we digitally band-pass filtered the raw LFPs using a zero phase delay filter (K. Harris and G. Buzsaki). The phase was calculated using a Hilbert transform, and a corresponding phase was assigned to each spike. We limited our analysis to units that fired at least 100 times over the period analyzed. Functional directionality was calculated based on the pairwise phase consistency (PPC) of VTA multiunit spikes assigned based on the corresponding phases of the 2-7 Hz filtered NAc LFP. We calculated PPC of VTA spikes that were shifted in 2.5 ms steps ± 80 ms to 2-7 Hz NAc signals. We compared the lags of the maximum PPC value to determine VTA-to-NAc directionality, where negative lags correspond to VTA-to-NAc directionality.

### Single-unit analyses

We identified significantly light-modulated units by bootstrapping. Laser ON and laser OFF spikes were combined, binned, and randomly shuffled 30,000 times. We considered units light-responsive if the observed laser OFF vs. laser ON firing rate difference was greater than 95% of the firing rate differences from the shuffled data.

To determine the effects of restraint effects on firing rate, average firing rate was compared between a 5-minute baseline period during which mice explored a familiar environment and restraint during the same recording session. For task-evoked single unit activity, spikes were binned in 10-ms bins and averaged across trials per neuron. CS+ evoked single unit activity was calculated as the average firing rate per neuron during CS+ presentation. Identification of putative VTA dopamine and VTA GABA neurons was adapted from Cohen *et al*^26^. Neurons were first filtered Neurons were first filtered based on whether they were significantly task-modulated by performing a one-way ANOVA of firing during a 1 second baseline period, during the 1 second CS+ presentation, and during the 0.5 second delay period. Significantly task-modulated units were then classified as putative VTA dopamine neurons if they had significantly higher reward-evoked firing in the 0-0.15 second period after reward retrieval relative to pre-CS+ baseline firing rate, as well as no difference between pre-CS+ baseline firing rate and delay period firing rate. Significantly task-modulated units were classified as putative VTA GABA neurons if they exhibited a higher firing rate during the delay period than the pre-CS+ baseline firing rate.

### Quantification and statistical analysis

Statistical analysis was performed in Graphpad Prism 7 or MATLAB. Two-tailed tests were used throughout. Data was tested for normality to determine whether to use parametric or non-parametric tests. For behavioral experiments, Wilcoxon signed-rank tests or paired t-test were used to assess the effects of stress and optogenetic manipulations within animals. For between-animal comparisons, Wilcoxon rank-sum tests or two-sample t-tests were used. For calculation of the efficacy of Arch and eNpHR on oscillation inhibition, Bonferroni-corrected one-sample *t*-tests were used to compare each time period to 100%. For all analyses, the alpha level was 0.05. Error bars and shaded error bands represent the standard error of the mean.

## Funding and Disclosure

This work was supported by a grant from the NIMH to A.Z.H (K08 MH109735), by a NARSAD grant from the Brain Behavior Research Foundation (AZH), and by the Hope for Depression Research Foundation (J.A.G., A.Z.H.). A.Z.H. is an advisory board member of Genetika+, incorporated.

## Acknowledgments

We would like to thank René Hen, Francis Lee, and Christian Lüscher for their feedback. We would also like to thank Mihir Topiwala, Joseph Stujenske and Ekaterina Likhtik for their early technical and analytic training as well as members of the Gordon and Hen labs for technical assistance and discussions. D.C.L. and A.Z.H. designed and analyzed the experiments. D.C.L., L.A.C, L.K, E.S.H., A.I.A., A.J.P., L.Y., Z.H.B, A.F. and A.Z.H. conducted the experiments and analyzed the data. J.A.G. assisted with the design, implementation, analysis and interpretation of experiments. D.C.L and A.Z.H. interpreted the results and wrote the paper.

## Data availability

The data that support the findings of this study are available from the corresponding author upon reasonable request.

## Code availability

The custom-written code used to analyze data is available from the corresponding author upon reasonable request.

## Notes

### Competing Interest Statement

AZH is an advisory board member of Genetika+, inc

