## Supplemental Figures for "Ventral tegmental area GABA neurons mediate stress-induced anhedonia"

#### Extended Data Figure 1

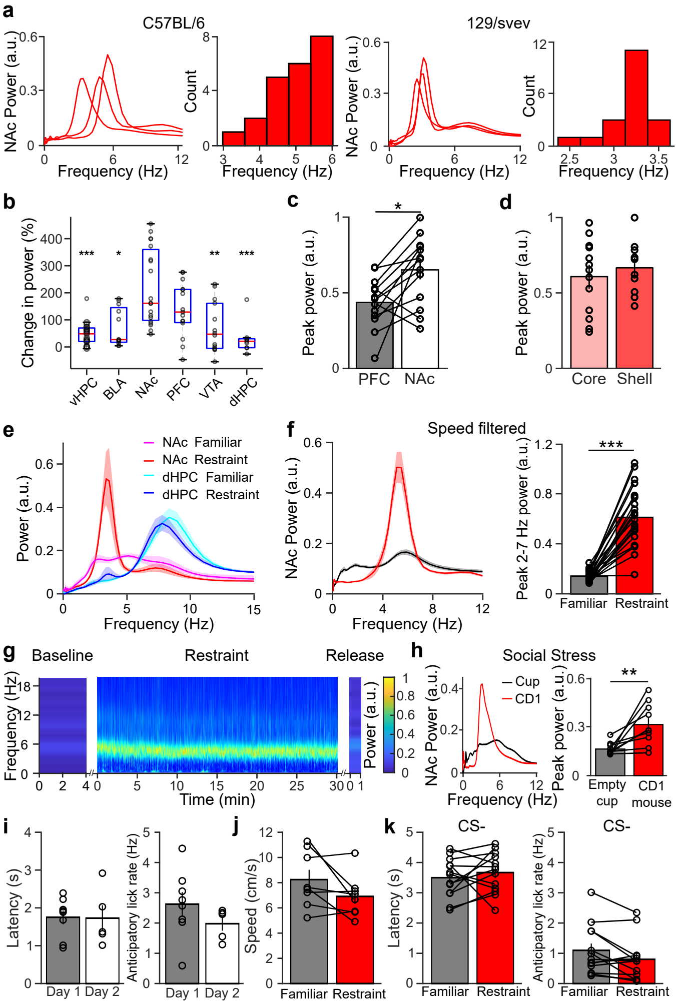

#### Extended Data Figure 2

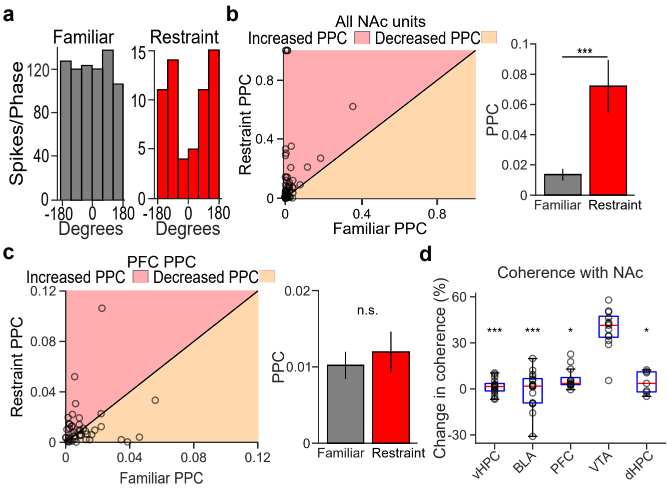

#### Extended Data Figure 3

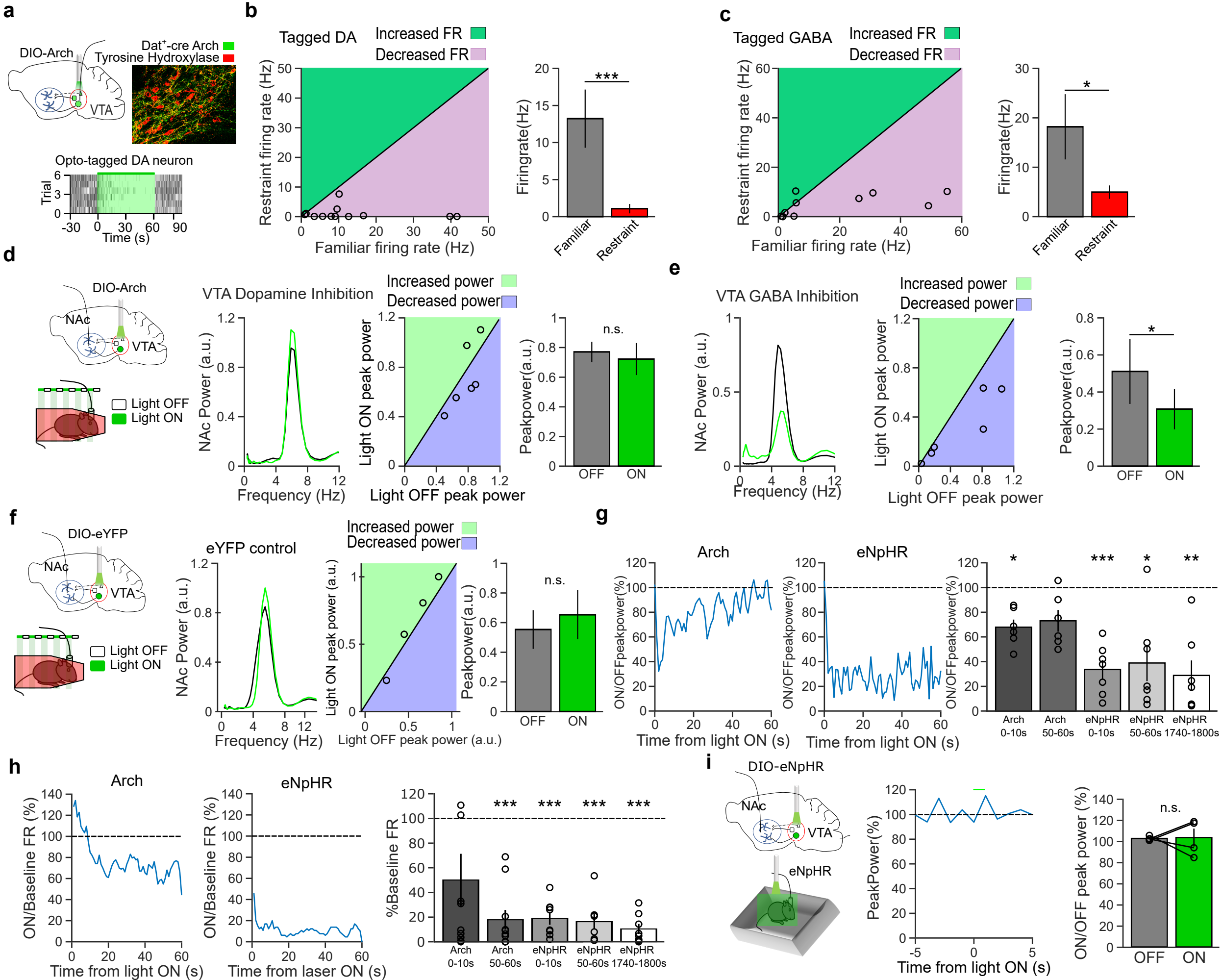

#### Extended Data Figure 4

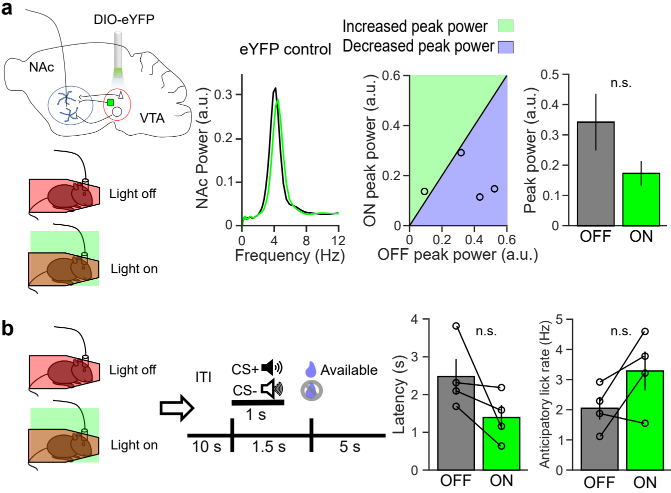

### Extended Data Figure 5

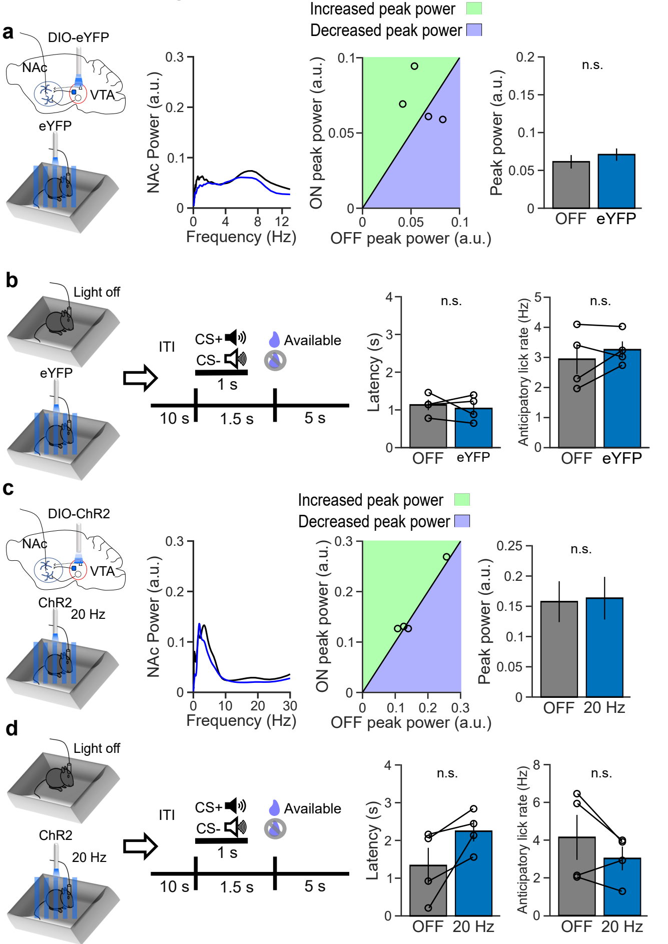
